# The representational dynamics of the animal appearance bias in human visual cortex are indicative of fast feedforward processing

**DOI:** 10.1101/2023.02.07.525897

**Authors:** Chiu-Yueh Chen, Gaëlle Leys, Stefania Bracci, Hans Op de Beeck

**Author notes:** Corresponding author at: Tiensestraat 102 - box 3714, 3000 Leuven, Belgium. E-mail address (H. Op de Beeck).

## Abstract

The human visual system has a seemingly unique tendency to interpret zoomorphic objects as animals, not as objects. This animal appearance bias is very strong in the ventral visual pathway as measured through functional magnetic resonance imaging (fMRI), but it is absent in feedforward deep convolutional neural networks. Here we investigate how this bias emerges over time by probing its representational dynamics through multivariate electroencephalography (EEG). The initially activated representations to lookalike zoomorphic objects are very similar to the representations activated by animal pictures and very different from the neural responses to regular objects. Neural responses that reflect the true identity of the zoomorphic objects as inanimate objects are weaker and appear later, as do effects of task context. The strong early emergence of an animal appearance bias strongly supports a feedforward explanation, indicating that lack of recurrence in deep neural networks is not an explanation for their failure to show this bias.

## 1. Introduction

The sensory systems of all animals have evolved and developed to detect the most important events in their environment. In many cases, those events are related to the presence of other animals. Prey animals have to detect predators, such as shown by the special sensitivity of rodent behavior and neural responses to looming objects that might signal the approach of a predator bird (Li et al., 2021; Yilmaz and Meister, 2013). The neural systems of predators are set up to detect and catch prey, illustrated by the existence of fly detectors in the frog’s brain (Barlow, 1953) and prey catching behavior in mice (Hoy et al., 2016). Many animals care about processing the behavior of conspecifics, resulting in elaborate processing of social stimuli (Powell et al., 2018; Sliwa and Freiwald, 2017).

In the human visual system, studies have revealed the existence of brain regions specialized for socially relevant stimuli such as faces and bodies (Downing et al., 2001; Kanwisher et al., 1997). These regions display a sensitivity for the degree of animacy, with a graded selectivity for how similar the face and body properties of a particular animal are to the human face and body (Ritchie et al., 2021). As a result, animacy comes out as a primary dimension characterizing object representations in human cortex and perception (Bracci and Op de Beeck, 2016; Kriegeskorte et al., 2008b; Mur et al., 2013; Yargholi and Op de Beeck, 2022). In addition to this general selectivity for animacy, human visual cortex also shows a bias to process non-animate or ambiguous stimuli as being animate. The term pareidolia is used for the general phenomenon of giving a meaningful interpretation to a random pattern or shape. Very often this interpretation is in terms of an animal form or face. Examples from daily life are numerous. We see all sorts of shapes, mostly animals, in clouds. We detect faces and human forms in rock formations and pizza. In the lab, participants interpret shape stimuli as complex animate forms even when performing simple and boring discrimination tasks (e.g., Op de Beeck et al., 2003; Op de Beeck, 2012). Perceived curvature might be a particularly important mid-level perceptual feature for this perception of animate forms (Long et al., 2017).

In some cases, the illusory perception of animacy dominates the overall processing of the presented objects in visual cortical processing, resulting in an animal appearance bias. Bracci et al. (2019) introduced a stimulus design with so-called zoomorphic or lookalike objects: non-animate objects that are made to look like an animal, such as a cow-shaped mug. It is still easy to interpret the lookalike objects for what they really are, objects, rather for what they appear to be, animals. When judging similarity, human observers considered the lookalike objects as somewhere in between animals and objects, a bit closer to inanimate objects than to animals. Feedforward deep neural networks (DNNs) exhibited the same behavior to an even greater extent, grouping the lookalike objects with inanimate objects. We could refer to this tendency as an object bias. Nevertheless, the neural response to these lookalike objects as measured through functional magnetic resonance imaging (fMRI) was almost indistinguishable from the response to actual animate objects, showing that human visual cortex is strongly affected by the appearance of the stimuli as animals. This result was even found when subjects were doing a task in which they had to group the lookalike objects with objects.

However, it is unclear how this animal appearance bias emerges during information processing. By using fMRI, Bracci et al. (2019) obtained a time-averaged view of representational similarity. Such data cannot distinguish different hypotheses about how representations evolve over time. A first possibility is that the animal appearance of the lookalike objects is detected early on in the first feedforward sweep of information processing. This would be consistent with the findings from a recent study on face pareidolia (Wardle et al., 2020). Stimuli that elicit face pareidolia are associated with an increased activity in face-selective brain regions and early face-selective electrophysiological responses (Wardle et al., 2020). Note though that despite the speed of processing of illusory faces, early electrophysiological responses to an illusory face were still more object-like than face-like, as was also the case for the (time-averaged) fMRI responses. In contrast, the animal bias in Bracci et al. (2019) was much larger than the face bias/pareidolia for the stimuli of Wardle et al. (2020), complicating the generalization between these two phenomena. Furthermore, the very stereotypical nature of face templates might speed up face detection relative to the detection of animal appearance, which is indeed supported by the earlier emergence of face clusters compared to animate/inanimate clusters in cortical representational spaces (Kietzmann et al., 2019). As a result of these differences, it is uncertain to which extent early feedforward processing would underlie the strong animal bias.

A second hypothesis about the emergence of the animal bias is that the processing of the animal-like appearance of lookalike objects might be present from the start but in addition increases over time. Such increase could depend on recurrent processing after the initial feedforward sweep of information processing. Recently there have been several reports that feedforward DNNs cannot fully capture the representational dynamics in human visual cortex (Kietzmann et al., 2019) and cannot explain human performance in difficult object recognition tasks (Seijdel et al., 2021; Tang et al., 2018). The very different behavior of feedforward DNNs (lookalikes processed as objects) and human visual cortex (lookalikes processed as animals) might be due to the fact that human visual cortex relies upon recurrent processing to process the animal-like appearance of these lookalikes. The gradual increase in the animacy representation might also explain why the animal bias measured by Bracci et al. (2019) is much stronger than face pareidolia effects (Wardle et al., 2020).

In the present study, we investigated the representational dynamics and task dependence of the animal bias for lookalike zoomorphic objects using electroencephalography and fMRI-EEG fusion. We find that the initially activated representations to lookalike objects are very similar to the representations activated by animal pictures. Neural responses that reflect the true identity of the lookalikes as inanimate objects are weaker and appear later. Task effects of the relevance of the animal appearance versus object identity were relatively minor and confined to later time points. In sum, the bias to process lookalike objects as if they are animals is particularly strong in the initial response to these lookalike objects.

## 2. Methods

### 2.1 Participants

30 healthy volunteers (23 females; mean age, 21 years) were recruited online through a university online recruitment system (SONA). The volunteers received either course credit or monetary rewards. Most volunteers were belonging to the student population of KU Leuven and there was no restriction in terms of gender. This study was approved by the KU Leuven Social and Societal Ethics Committee (G-2020-2379). All participant’s data were organized according to the brain imaging data structure (BIDS) (Pernet et al., 2019).

### 2.2 Stimuli and experimental design

Stimuli consisted of nine triads, resulting in a total of 27 stimuli (see Figure 1). Within these triplets, visual appearance and category information (animacy) were manipulated. Each triplet consists of one animate, one inanimate, and one zoomorphic object that looks like the animal and is matched to the object (e.g., a cow, a mug, and a cow-shaped mug). All of the images were gray-scaled and used before by Bracci et al. (2019).

**Figure 1.**
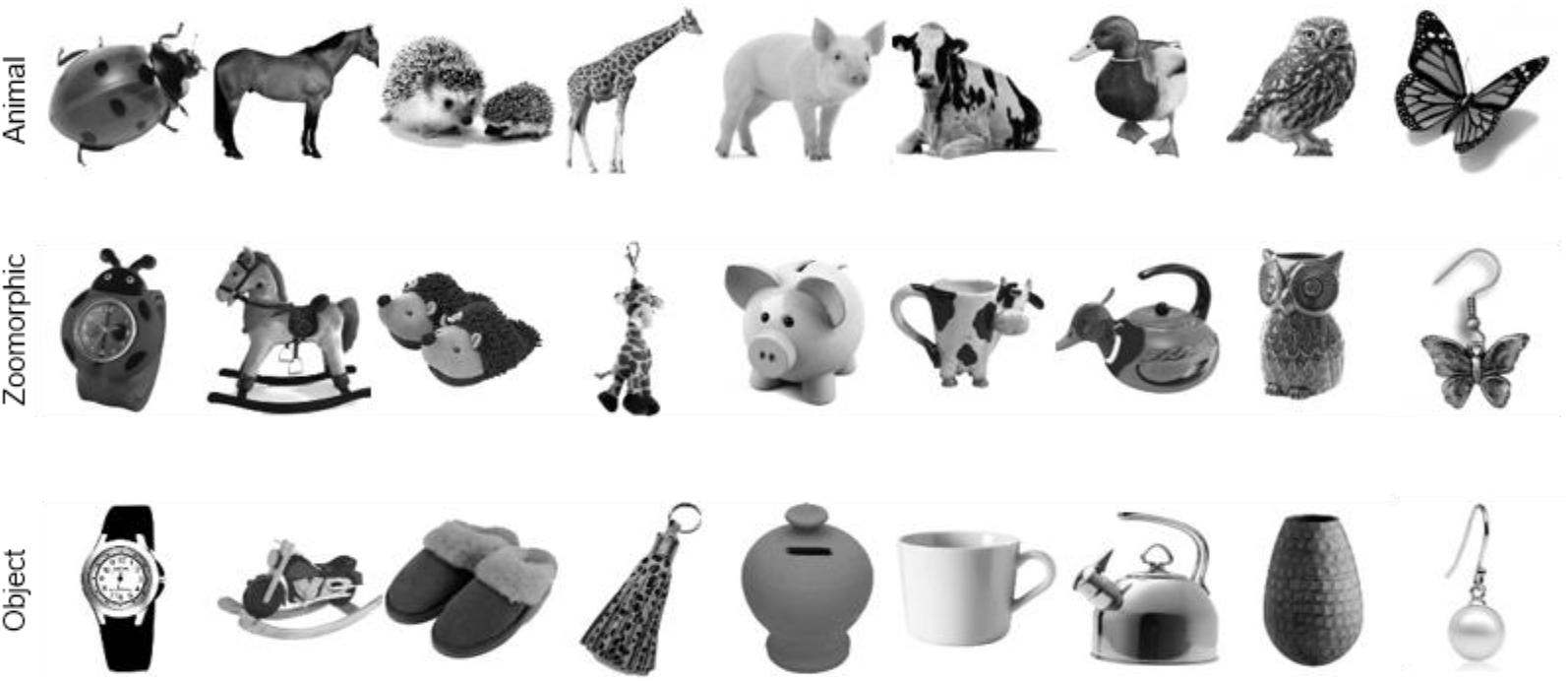
Experimental stimuli. The stimulus set consisted of 27 stimuli in three categories (animate, zoomorphic, and inanimate objects) and was used in the study of Bracci et al. (2019).

Participants performed two tasks: an animacy task and an animal appearance task. In the animacy task, participants judged animacy (“Does this image depict a living animal or an object?”). Participants responded ‘object’ to the objects as well as to the zoomorphic objects. In terms of response tendency for the zoomorphic objects, this can be rephrased as an object bias (lookalikes classified with the objects). In the appearance task, participants judged animal appearance (“Does this image look like an animal?”). Participants responded ‘animal’ to the animals as well as to the zoomorphic objects, which we refer to as an animal (appearance) bias. Participants completed seven runs of 108 trials on each task. Within each trial, an image was presented for 500 ms in the center of the screen, with a random inter-trial interval ranging between 700 ms to 1100 ms.

### 2.3 EEG recording and preprocessing

EEG signals were recorded with a 128-channel active electrode system arranged according to the extended radial system, using an ActiveTwo amplifier at a sampling rate of 1024 Hz (BioSemi, Amsterdam, Netherlands). A photosensor tracked the exact onset time of each stimulus by detecting a concurrent change from black to white in a corner of the screen.

Participants were around 60 cm away from a BenQ XL2411 screen (24 inches, 60 Hz, resolution of 1920 × 1080 pixels), in a dark room. The stimulus presentation was controlled using a script constructed with the PsychoPy experiment builder (Peirce et al., 2019). All stimuli were presented at a resolution of 324 × 324 pixels.

Offline preprocessing was conducted using the FieldTrip toolbox (Oostenveld et al., 2011) in MATLAB R2020b. To reduce the slow drift noise and the power line noise, a 2 Hz high-pass filter and a 50 Hz notch filter were used. Traces were demeaned per run (baseline correction), referenced to the average of all 128 channels, and then resampled to 250 Hz.

To remove artifacts for eye movements, muscle, heartbeats, and the channels containing excessive noise, independent component analysis (ICA) was performed using EEGLAB (Delorme and Makeig, 2004). Subsequently, the components were labeled and removed using ICLabel (Pion-Tonachini et al., 2019). Afterwards, the artifact-free data was segmented into 700 ms epochs from -200 ms to 500 ms relative to stimulus onset. We define stimulus onset relatively to the onset of the photosensor, which is later than the time at which the stimulus presentation script gives the command to flip the screen.

### 2.4 Category-level decoding analysis

To determine the amount of object category information contained in EEG data, decoding analyses were applied (for review, see Grootswagers et al., 2017). A temporal searchlight analysis using linear discriminant analysis (LDA) classifier was performed, as implemented in the CosMoMVPA toolbox (Oosterhof et al., 2016). We implemented this searchlight analysis nine times per participant, combining the three pairwise contrasts of the three categorical distinctions (animal; lookalike; object) with three task circumstances: both tasks together (combining data from 14 runs), and each task separately (animacy task and appearance task). The temporal searchlight analysis included the multi-sensor signal from all 128 sensors. The temporal neighborhood consisted of each center time point with four neighboring time points, moving across all time points from -200 ms to 500 ms. The LDA classifier was trained and tested using leaving-one-run-out cross-validation.

### 2.5 Scalp topography of category-level decoding

A spatiotemporal searchlight analysis was performed with a different spatial neighborhood setting in the sensor space. For each sensor, the sensor and its nine nearest neighboring sensors in the configuration from Biosemi 128 electrode cap formed a neighbourhood. After iterating across all 128 sensors and across all time points, and averaging across all participants, the resulting maps yield the topography of category-level decoding accuracy.

### 2.6 Representational similarity analysis

Representational similarity analysis was used to evaluate the similarity for all individual image pairs over time. We again used temporal searchlight analyses including multi-sensor patterns from all sensors and each time point plus four neighboring time points. For each participant and each task, the LDA classifier was trained and tested to discriminate between each pair of individual images (27×26 pairs), using leaving-one-run-out cross-validation. The pairwise image decoding accuracy was used as a measure of neural dissimilarity. As a result, we obtain a neural dissimilarity matrix for each time point.

The dissimilarity matrices were compared with other data modalities in so-called representational similarity analyses (Kriegeskorte et al., 2008a). Only the upper half of the matrices were used in these analyses. First, the dissimilarity matrices were correlated with the predictions from two conceptual models, an animacy model and an appearance model (Bracci et al., 2019). For the animacy model, the lookalikes were expected to evoke similar activation patterns as inanimate objects, in accordance with an object bias. For the appearance model, the lookalikes were expected to evoke similar activation patterns as animate objects, showing an animal bias. Second, we performed fMRI-EEG fusion. We used the fMRI dissimilarity matrices from the three regions of interest in Bracci et al. (2019): early visual cortex (EVC), posterior ventro-temporal cortex (pVTC), and anterior ventro-temporal cortex (aVTC). The fMRI matrices are averaged across 16 participants.

### 2.7 Statistical inference

Statistical significance was assessed using the threshold-free cluster enhancement procedure (TFCE) (Smith and Nichols, 2009) and multiple-comparison correction with null distributions created from 1000 bootstrapping iterations, all as implemented in the CoSMoMVPA toolbox. For category-level decoding and the decoding difference between category pairs, the null hypothesis of no difference was conducted by a permutation test that shuffled the category labels on each participant 100 times. For individual image pair decoding, the null hypothesis of chance (50%) was set. For correlation between neural dissimilarity matrices and models and their differences, the null hypothesis of zero correlation was used. The threshold was set at z > 1.96 and z < -1.96 (i.e., TFCE corrected p < 0.05, two-tailed).

### 2.8 Data and code availability statement

Stimulus presentation code, analysis code, and data will be made available through the Open Science Framework at xxx.

## 3. Results

### 3.1 How are lookalikes processed relative to animals and objects?

We grouped the 27 images in three category-level conditions: Animate (animals), lookalike and inanimate (regular objects). We trained linear classifiers for the three possible pairwise contrasts between these three conditions using all trials from all runs but one, and tested these classifiers on the individual trials of the left-out run. This procedure was iterated until all runs served as left-out run. Fig. 2 shows the resulting test performance averaged across all participants, taking all runs together (panel A), or separately for the two task contexts (panels B-C).

**Fig. 2.**
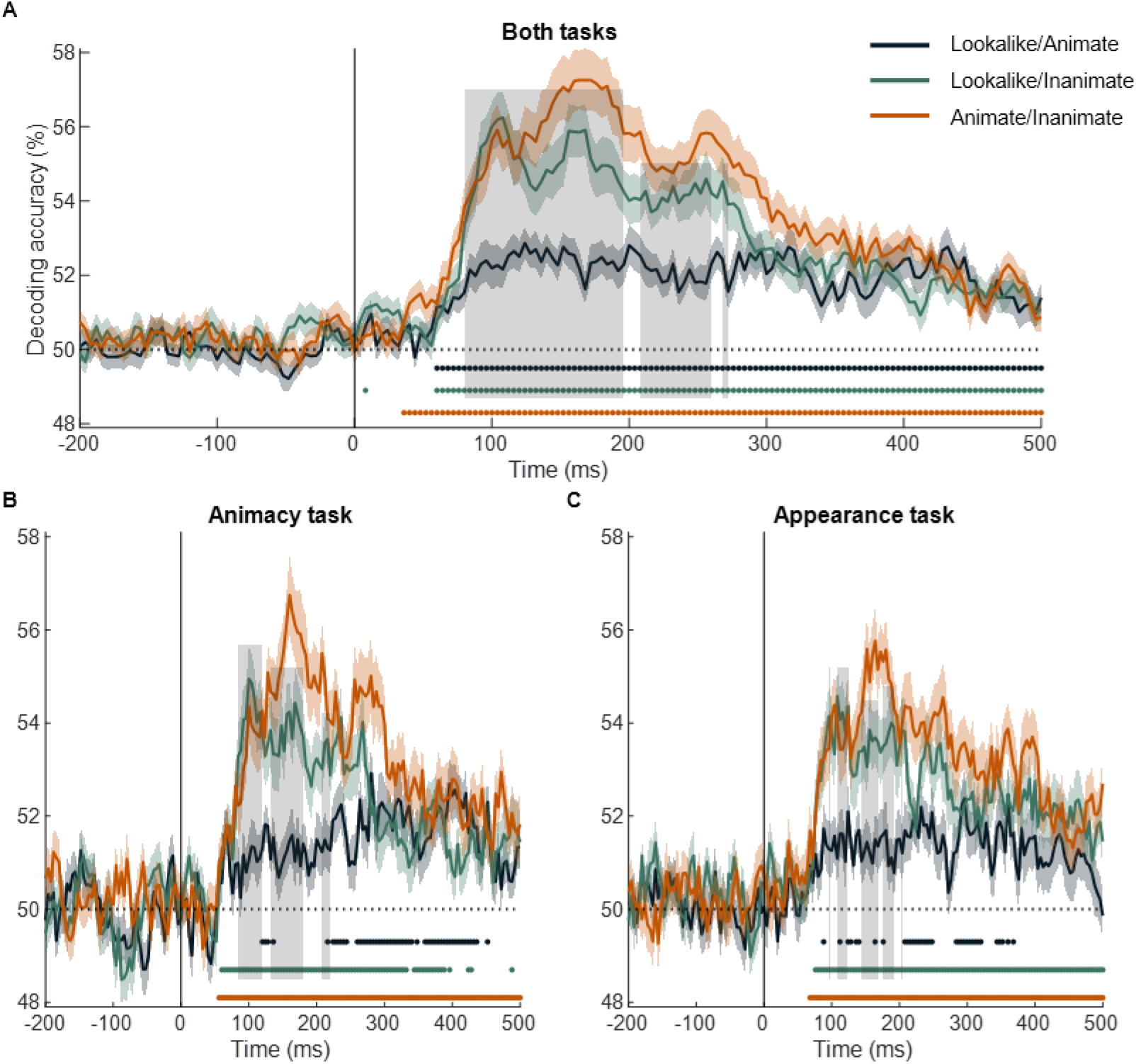
Time course of decoding for condition pairs. The condition-wise decoding accuracies over time are shown for the data combined across the two tasks (A), for the animacy task (B), and for the appearance task (C). Different lines show the decoding performance for each condition pair: animate against lookalike (blue), lookalike against inanimate (green), and animate against inanimate (orange). The light-coloured regions above and below the mean lines indicate standard error across subjects (n = 30). Marks above the x axis indicate time points where the decoding performance is significantly greater than chance. The vertical gray lines indicate the time points where the decoding difference between the lookalike and inanimate (green), and lookalike and animate (blue) is significantly different than zero.

We first analyzed the data combined across the two behavioral tasks. The decoding across time for the distinction between animate and inanimate serves as a benchmark for the other distinctions (Fig. 2A, orange line). This decoding goes up towards a first peak around 104 ms, increases further to a second peak around 160 ms, and then gradually decreases towards the end of stimulus presentation but remains significant throughout.

We find almost equally high decoding for the distinction between lookalike and inanimate (Fig. 2A, green line). The initial peak around 108 ms virtually has the same height as for the distinction between animate and inanimate. Afterwards decoding performance declines but remains high. In comparison, the decoding of lookalikes versus animate is much lower (Fig. 2A, blue line), significantly lower throughout most of the interval from 80 to 272 ms (grey area in figures). Summarized, the pairwise decoding of the three conditions suggests that lookalike objects are mostly represented as if they are animals for the first hundreds of milliseconds of the neural response.

The temporal dynamics and relative decoding strengths were very similar in the two task settings, the animacy task and the appearance task (Fig. 2B-C). This is most striking for the animacy task. In this task, subjects are asked to group the lookalikes with the inanimate objects. Nevertheless, lookalikes are represented as most similar (lowest decoding) to the animals throughout the early part of the neural response. In later parts of the response, the decoding of lookalikes is similar for the two contrast conditions, versus animate and versus inanimate, and this is found in each task context.

We performed a spatiotemporal searchlight analysis to explore which electrode neighbourhoods would provide the strongest signal to distinguish the three stimulus conditions in a pairwise manner. EEG is not well suited for anatomical localization, but as far as there are differences between sensors in how much they allow for category-level decoding we would expect these sensors to be over occipitotemporal cortex where these categorical differences are typically reported in fMRI. Indeed, for each category-level distinction, we found the highest decoding around occipitotemporal electrodes, maybe with a slight bias towards the right (Fig. 3). In addition, these topographic plots further confirm the overall differences in effect size between distinctions, with larger effects for animate versus inanimate and for lookalike versus inanimate than for lookalike versus animate. However, the topographies do not suggest clear differences in the spatial distribution of the diagnostic signals between the three pairwise comparisons. Nor is there an obvious change in this topography across time beyond the expected reduction in amplitude towards later time points.

**Fig. 3.**
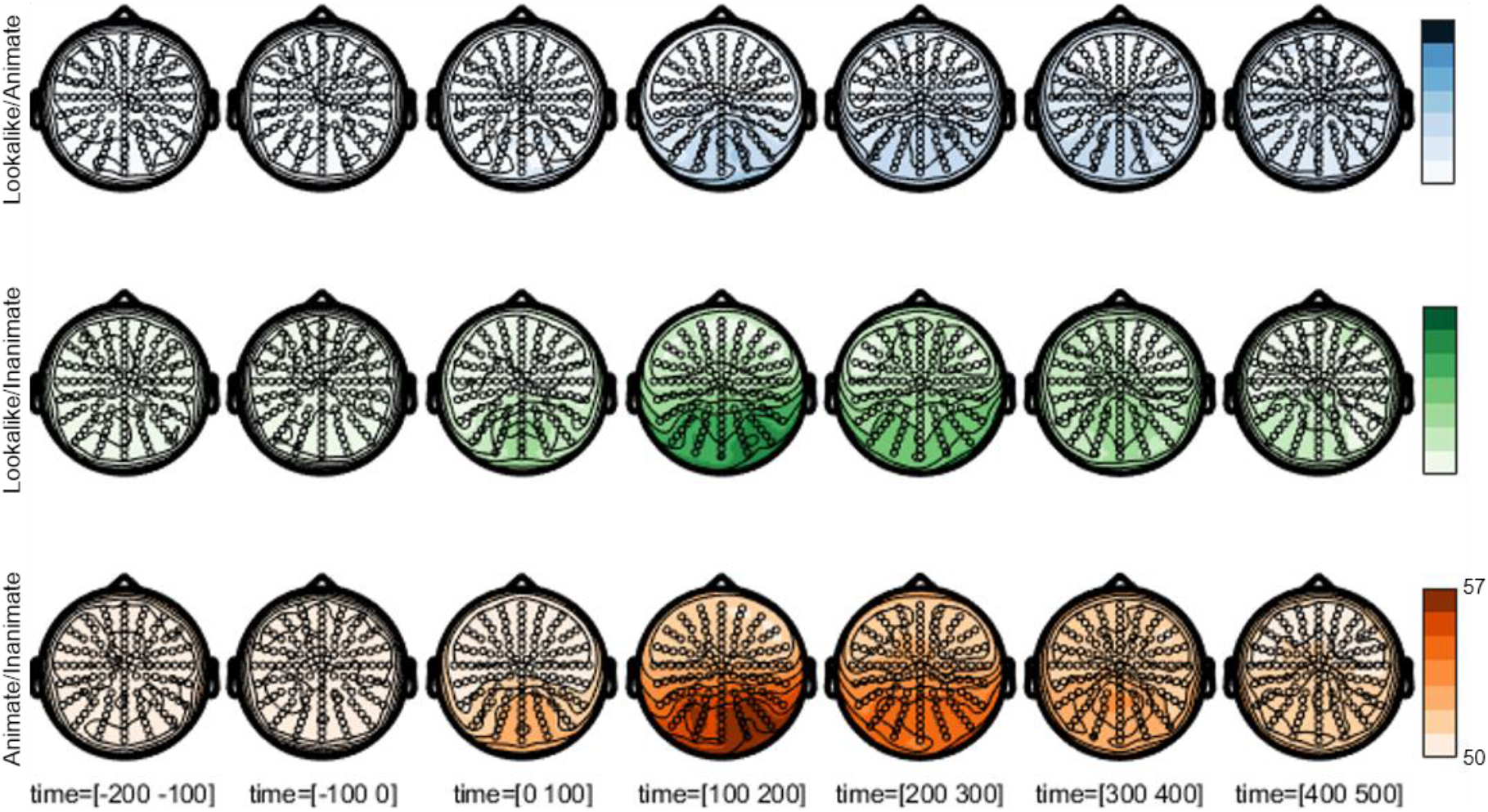
Scalp topography of decoding for condition pairs. Scalp plots show the topography of average decoding performance in seven 100 ms time intervals for each condition pair: lookalike and animate (blue), lookalike and inanimate (green), and animate and inanimate (orange).

### 3.2 Representational similarity in pairwise image differences

When we train classifiers to decode the differences between the three aforementioned stimulus conditions, we lose information about differences among individual images. This grouping at the level of conditions also increases the challenge for the classification, as the classifier can only use features that are common to the images in a condition. With this in mind, it might not be a surprise that the peak decoding performance is higher when we classify individual pairs of images, despite the fact that this classification is based upon much less training data. This peak decoding of individual image classification reaches up to 60% (Fig. 4A), up from around 57% in the condition-wise decoding (Fig. 2). The curve as a function of time now shows a more prominent early peak, probably because decoding can be based upon more simple features that distinguish individual images. The curve has a very similar shape in the different task contexts (Fig. 4B-C).

**Fig. 4.**
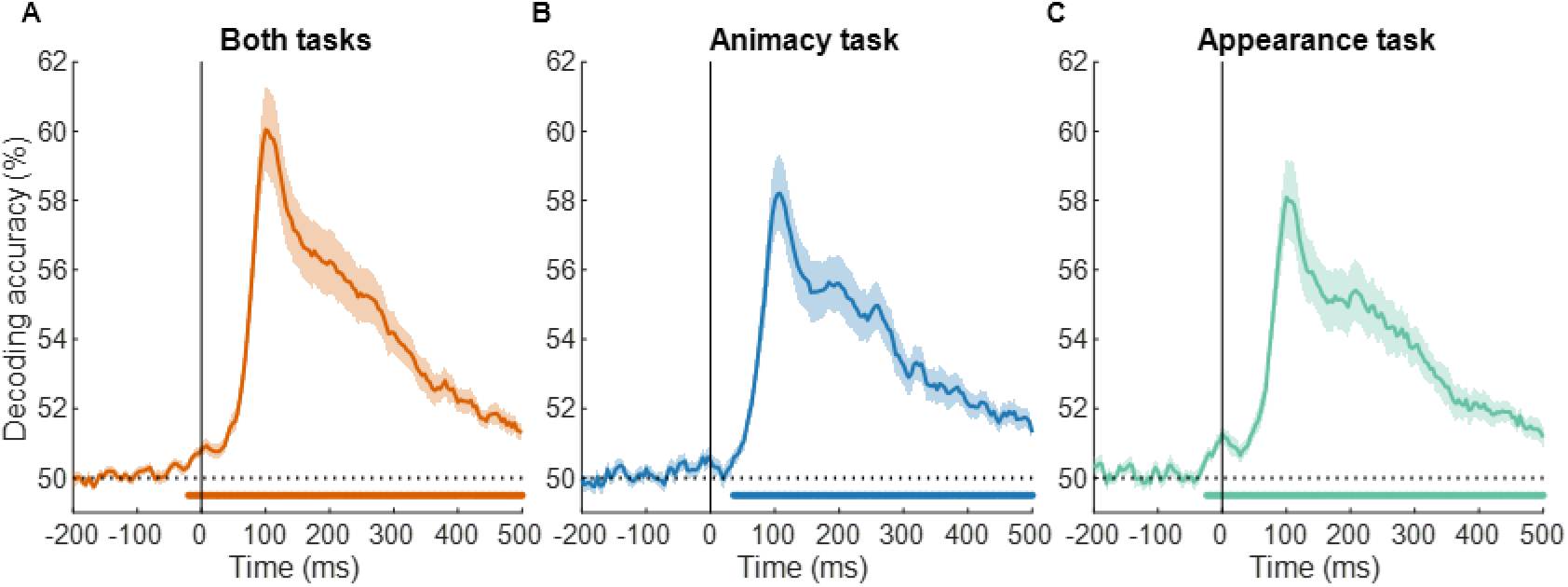
Time course of decoding for image pairs. The pairwise decoding accuracies over time, averaged across all 27×26 stimulus pairs, are shown for the data combined across the two tasks (A), for the animacy task (B), and for the appearance task (C). The shaded regions above and below the mean lines indicate standard error across subjects (n=30). Marks above the x axis indicate the time points where decoding performance is significantly different from chance.

The overall decoding hides the truly interesting level of analysis, which is the representational dissimilarity matrix showing the decoding of all individual image pairs (Fig. 5). We have one such matrix for each time point. The supplemental information shows all time points as a movie, in Fig 5 we display the matrices at four time points: 0 ms, 80 ms, 100 ms, and 168 ms. We show the matrices for both tasks analyzed together and for the two tasks separately. The task-specific figures reveal the replicability of the matrices across tasks. In all tasks, we find a blue matrix (overall decoding close to chance) around 0 ms, which transforms into a green-yellow matrix around 100 ms. Around time 160 ms there is an obvious quadrant structure with a large quadrant in the top left which relates to a clustering of lookalikes with animals.

**Fig. 5.**
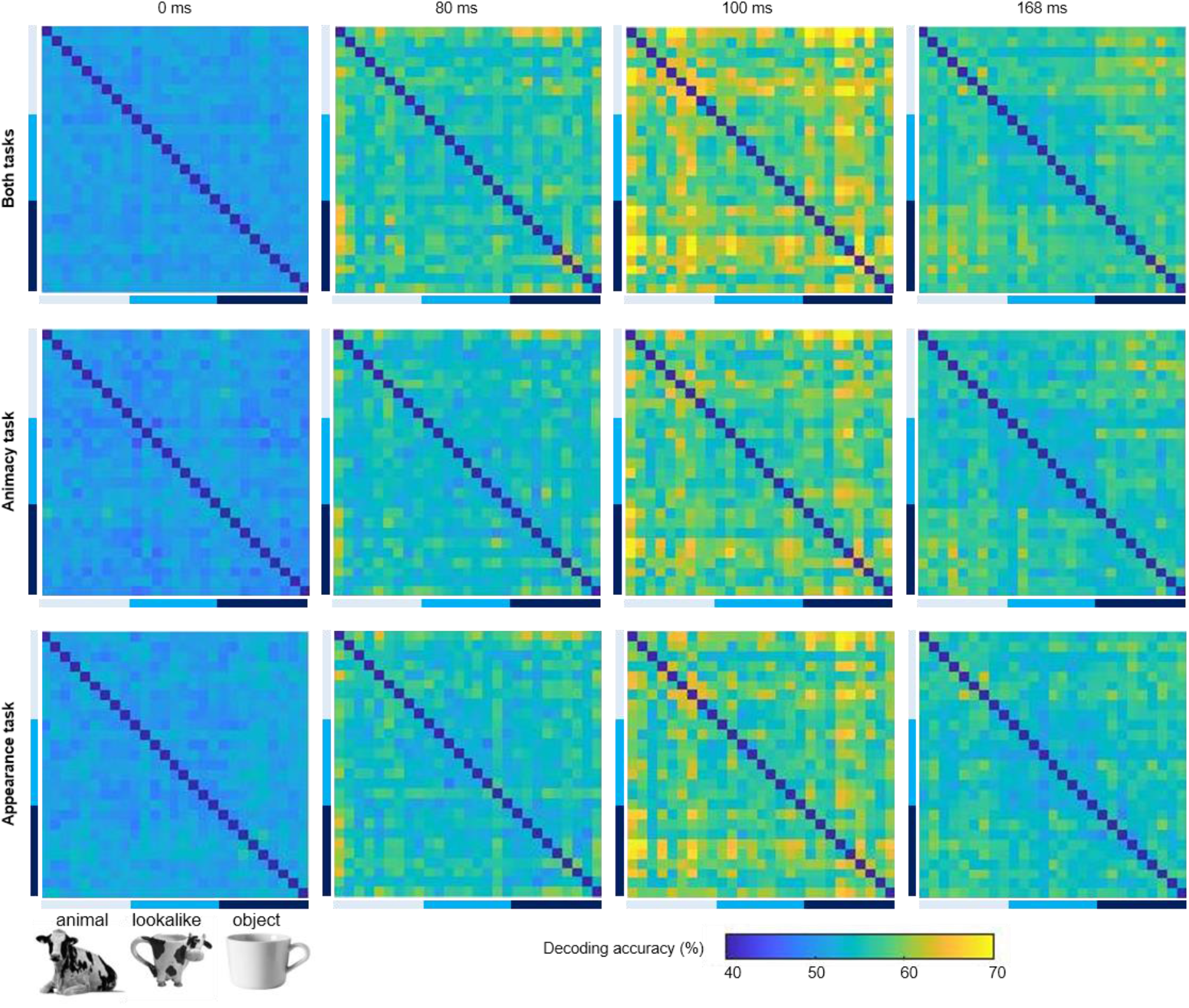
The decoding accuracies for all image pairs at 0 ms, 80 ms, 100 ms, and 168 ms. The pairwise decoding matrices at selected time points are shown for the data combined across the two tasks, for the animacy task, and for the appearance task. The higher decoding accuracy (yellow) corresponds to greater neural dissimilarity. Note that there are no data values along the diagonal, these values are pre-set at the bottom end of the colour scale.

To analyze this pattern more quantitatively, we correlated the matrices at each time point with the matrices from the two a priori conceptual models: the appearance model, in which lookalikes are clustered with animals, and the animacy model, in which lookalikes are clustered with the inanimate objects. Taking the data from both task settings together, we find a significant correlation with the appearance model throughout a long-time interval, but much less so with the animacy model (Fig. 6A). For part of the time, most prominently around 160 -200 ms, the correlation with the appearance model is significantly higher than with the animacy model.

**Fig. 6.**
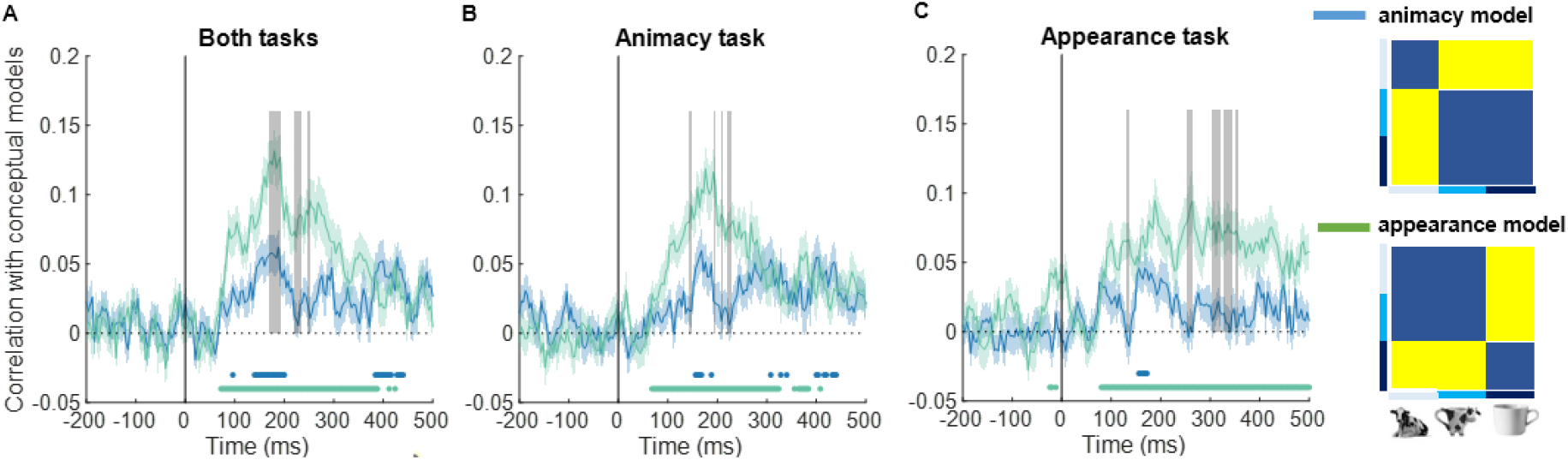
Correlation between decoding accuracy and theoretical models over time. Time course of correlation between decoding accuracy and theoretical models is shown for the data combined across the two tasks (A), for the animacy task (B), and for the appearance task (C). The shaded regions above and below the mean lines indicate standard error across subjects (n = 30) and the marks above the x axis indicate time points where the correlation is significantly different than zero. The vertical gray lines indicate time points where the difference between the animacy model (blue) and the appearance model (green) is significantly different than zero.

Through these correlations with model matrices, we also obtain a first clear effect of task setting. In the animacy task (Fig. 6B), the later part of the responses showed virtually identical correlations with the two models. In contrast, in the appearance task, there was a significantly stronger correlation with the appearance model than with the animacy model (Fig. 6C).

### 3.3 FMRI-EEG fusion

Bracci et al. (2019) investigated the representational similarity with the same stimulus design with fMRI. fMRI provides time-averaged data with sufficient spatial resolution to distinguish between separate brain areas. The dissimilarity matrices at the bottom of Figure 7 represent their findings for three separate ROIs: early visual cortex (EVC), posterior ventro-temporal cortex (pVTC), and anterior ventro-temporal cortex (aVTC). Bracci et al. (2019) found that the fMRI similarity patterns in EVC correlated with neither model, while pVTC and aVTC showed a significantly stronger correlation with the appearance model.

**Fig. 7.**
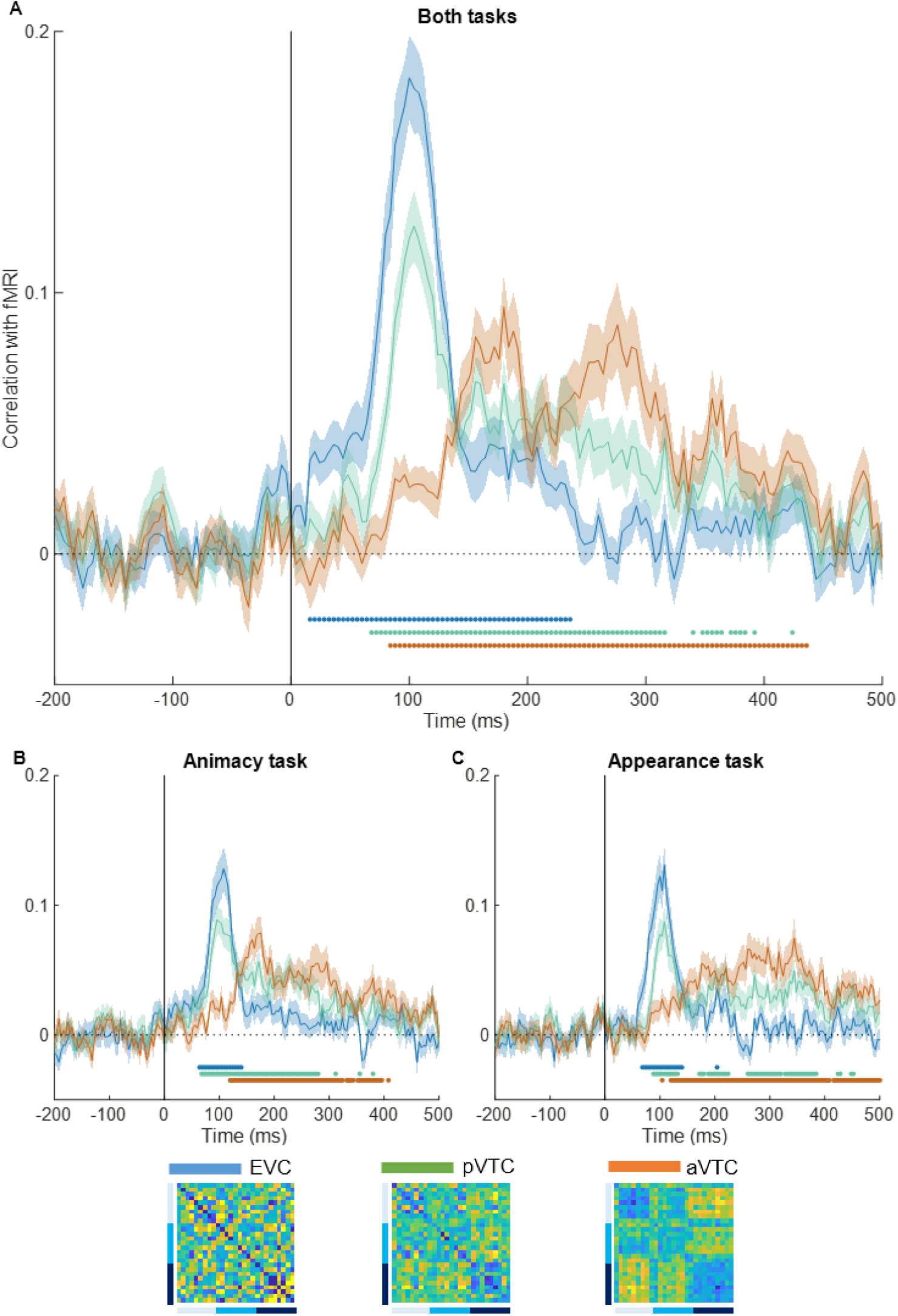
fMRI-EEG fusion. Time courses of correlation between the EEG pairwise decoding matrices and fMRI similarity patterns are shown for the data combined across the two tasks (A), for the animacy task (B), and for the appearance task (C). The shaded regions above and below the mean lines indicate standard error across subjects (n = 30) and the marks above the x axis indicate time points where the correlation is significantly different from zero.

Here we report how the time-averaged activity in these three brain regions is related to the temporal dynamics as measured through EEG (Fig. 7A). Consistent with our knowledge of the visual processing hierarchy, the representational similarity matrix in EVC correlated strongly with relatively early time points in the EEG (peak: 100 ms), pVTC still showed a clear early peak but with some sustained correlations, and aVTC correlations were initially very weak and increased towards a peak at much later time points (first peak around 156 ms). The temporal development was qualitatively similar in the two task conditions (Fig. 7B-C).

These findings reveal that the early bias to represent lookalikes as animals in EEG data cannot be due to aVTC only, as this region shows almost no correlation with EEG in these early time points. It is interesting to visually compare the EEG condition-wise decoding (Fig. 2) and representational similarity analyses (Fig. 6) with the fMRI-EEG fusion (Fig. 7). The temporal profile in the first two, summarized as a clear decoding early on but with a comparable peak later on, might be explainable by a combination of the pVTC and aVTC results in the fMRI-EEG fusion.

## 4. Discussion

We investigated the representational dynamics underlying the animal appearance bias in the visual cortical processing for zoomorphic objects. The current findings demonstrate that the animal appearance bias emerges early on in the first feedforward sweep of information processing. Early cortical responses tend to process lookalike objects more like animals than like regular, inanimate objects. A first line of evidence is the very strong early decoding of the distinction between lookalike objects and regular inanimate objects, combined with a much weaker decoding of the distinction between lookalike objects and animals. A second line of evidence is a strong early correlation with the so-called appearance model in which lookalike objects are clustered with animals. These effects persist for at least 200 ms, after which the animal appearance bias fades out. In the later responses it depends a bit which task participants are performing. In particular, the stronger correlation with the appearance model persists into later time points only when participants are performing an appearance task in which they group the lookalikes with animals.

There is little room for doubt with these data. With 30 participants, we obtained very reliable results. Across both tasks together and replicated in each task context separately, we find that early responses are very much biased towards differentiating lookalike objects from inanimate objects, more so than differentiating lookalike objects from animals. The strength of this animal appearance bias, its relatively long duration, and its resilience to task effects is overall consistent with earlier findings obtained with fMRI (Bracci et al., 2019). The most important new piece of information here is that this bias is present already in the initial responses, and it is particularly strong early on. Only towards much later time points the lookalikes seem to fall more in the middle between animals and inanimate objects. Only in these later time points, there is some indication of an effect of task context. The spatiotemporal searchlight analyses suggest similar topographies of the diagnostic neural signals across pairwise comparisons and across time, so we have no strong indications that different brain regions would be involved. Broadly speaking, and keeping the low spatial resolution of EEG in mind, the different categorical distinctions and the decoding at different time points seem to be supported by the same set of regions (see e.g., Graumann et al., 2022).

The strong early emergence of an animal appearance bias strongly supports the first hypothesis that the processing of the animal appearance in lookalike objects is carried out in the initial feedforward sweeps of information processing. This outcome is consistent with the findings of Wardle et al. (2020) on face pareidolia. Note however that the strength of the effects is very different. In the case of Wardle et al., the object images that induced face pareidolia were still mostly processed as objects. The early face-like responses are small in comparison. Translating their findings to our design, what they found would be as if the lookalikes versus inanimate would have the lowest pairwise decoding, and lookalikes versus animate much higher but lower than inanimate versus animate in the initial part of the response. The pareidolia effect is small in such a case. The animal appearance bias as we see it in our design is much larger.

A close comparison with Wardle et al’s data also reveals another discrepancy: our onsets are faster. In our analyses, a first peak of decoding and correlations is found around 100 ms, while in Wardle et al. decoding only starts going up around 100 ms with much later peaks. Note that EEG/MEG articles typically do not mention how they define stimulus onset, and probably in the large majority of papers this is the time at which the presentation software gives the command to show the stimulus on the screen. This practice stands in contrast to the more precise benchmark that is often used in animal physiology studies where zero time is the time when the stimulus actually comes on the screen, as for example verified through a photodiode placed on the screen. We used the latter practice, and the two methods provide a (constant) 32 ms difference in stimulus onset in our setup. With our method we will have briefer but arguably also more precise latencies.

While our findings are consistent with the first hypothesis in terms of feedforward processing, they contradict an explanation in terms of recurrent processing. From this perspective the early emergence of the animal appearance bias further deepens the mystery of why deep neural networks do not show this animal appearance bias. It was an obvious way out to point to the absence of recurrent processing in these artificial networks, but this argument is no longer valid now that we know that the animal appearance bias in human visual cortex emerges in the initial feedforward sweep of information processing. Furthermore, a gradual build-up of the animal appearance bias over time as recurrent processing proceeds was also a possible explanation for why the animal appearance bias is so much stronger compared to for example face pareidolia.

The current findings will be important for constraining further computational studies aimed at understanding why the human visual system shows such a strong animal appearance bias and why it is already so prominent in the early feedforward processing of objects. One potential avenue is the implementation of a variety of training regimes that have been shown to change information processing in feedforward deep convolutional neural networks.

## Acknowledgements

This work was supported by the KU Leuven Research Council [grant number ZKD1090, C14/21/04]; and Fonds voor Wetenschappelijk Onderzoek FWO-Flanders [grant number G0D3322N]. CYC was supported by a KU Leuven–Taiwan doctoral fellowship; and the 2022 National Science and Technology Council Taiwanese Overseas Pioneers Grants (TOP Grants) for PhD Candidates.

## References

Barlow, H.B., 1953. Summation and inhibition in the frog’s retina. J Physiol 119, 69–88. https://doi.org/10.1113/jphysiol.1953.sp004829

Bracci, S., Op de Beeck, H.P., 2016. Dissociations and associations between shape and category representations in the two visual pathways. Journal of Neuroscience 36, 432–444. https://doi.org/10.1523/JNEUROSCI.2314-15.2016

Bracci, S., Ritchie, J.B., Kalfas, I., Op de Beeck, H.P., 2019. The Ventral Visual Pathway Represents Animal Appearance over Animacy, Unlike Human Behavior and Deep Neural Networks. The Journal of Neuroscience 39, 6513. https://doi.org/10.1523/JNEUROSCI.1714-18.2019

Delorme, A., Makeig, S., 2004. EEGLAB: an open source toolbox for analysis of single-trial EEG dynamics including independent component analysis. Journal of Neuroscience Methods 134, 9–21. https://doi.org/10.1016/j.jneumeth.2003.10.009

Downing, P.E., Jiang, Y., Shuman, M., Kanwisher, N., 2001. A Cortical Area Selective for Visual Processing of the Human Body. Science 293, 2470–2473. https://doi.org/10.1126/science.1063414

Graumann, M., Ciuffi, C., Dwivedi, K., Roig, G., Cichy, R.M., 2022. The spatiotemporal neural dynamics of object location representations in the human brain. Nature Human Behaviour 6, 796–811. https://doi.org/10.1038/s41562-022-01302-0

Grootswagers, T., Wardle, S.G., Carlson, T.A., 2017. Decoding dynamic brain patterns from evoked responses: A tutorial on multivariate pattern analysis applied to time series neuroimaging data. Journal of Cognitive Neuroscience. https://doi.org/10.1162/jocn_a_01068

Hoy, J.L., Yavorska, I., Wehr, M., Niell, C.M., 2016. Vision Drives Accurate Approach Behavior during Prey Capture in Laboratory Mice. Current Biology 26, 3046–3052. https://doi.org/10.1016/j.cub.2016.09.009

Kanwisher, N., McDermott, J., Chun, M.M., 1997. The Fusiform Face Area: A Module in Human Extrastriate Cortex Specialized for Face Perception. J. Neurosci. 17, 4302–4311. https://doi.org/10.1523/JNEUROSCI.17-11-04302.1997

Kietzmann, T.C., Spoerer, C.J., Sörensen, L.K.A., Cichy, R.M., Hauk, O., Kriegeskorte, N., 2019. Recurrence is required to capture the representational dynamics of the human visual system. Proceedings of the National Academy of Sciences of the United States of America 116, 21854–21863. https://doi.org/10.1073/pnas.1905544116

Kriegeskorte, N., Mur, M., Bandettini, P., 2008a. Representational Similarity Analysis – Connecting the Branches of Systems Neuroscience. Front Syst Neurosci 2, 4. https://doi.org/10.3389/neuro.06.004.2008

Kriegeskorte, N., Mur, M., Ruff, D.A., Kiani, R., Bodurka, J., Esteky, H., Tanaka, K., Bandettini, P.A., 2008b. Matching categorical object representations in inferior temporal cortex of man and monkey. Neuron 60, 1126–1141. https://doi.org/10.1016/J.NEURON.2008.10.043

Li, Z., Wei, J.-X., Zhang, G.-W., Huang, J.J., Zingg, B., Wang, X., Tao, H.W., Zhang, L.I., 2021. Corticostriatal control of defense behavior in mice induced by auditory looming cues. Nat Commun 12, 1040. https://doi.org/10.1038/s41467-021-21248-7

Long, B., Störmer, V.S., Alvarez, G.A., 2017. Mid-level perceptual features contain early cues to animacy. Journal of vision 17. https://doi.org/10.1167/17.6.20

Mur, M., Meys, M., Bodurka, J., Goebel, R., Bandettini, P.A., Kriegeskorte, N., 2013. Human object-similarity judgments reflect and transcend the primate-IT object representation. Frontiers in Psychology 4, 128. https://doi.org/10.3389/FPSYG.2013.00128/BIBTEX

Oostenveld, R., Fries, P., Maris, E., Schoffelen, J.-M., 2011. FieldTrip: Open Source Software for Advanced Analysis of MEG, EEG, and Invasive Electrophysiological Data. Computational Intelligence and Neuroscience 2011, 156869. https://doi.org/10.1155/2011/156869

Oosterhof, N.N., Connolly, A.C., Haxby, J.V., 2016. CoSMoMVPA: Multi-Modal Multivariate Pattern Analysis of Neuroimaging Data in Matlab/GNU Octave. Frontiers in Neuroinformatics 10, 27. https://doi.org/10.3389/fninf.2016.00027

Op de Beeck, H.P., 2012. The Distributed Nature of Visual Object Learning, in: Plasticity in Sensory Systems. Cambridge University Press, pp. 9–32. https://doi.org/10.1017/CBO9781139136907.002

Op de Beeck, H.P., Wagemans, J., Vogels, R., 2003. The effect of category learning on the representation of shape: dimensions can be biased but not differentiated. Journal of experimental psychology. General 132, 491–511. https://doi.org/10.1037/0096-3445.132.4.491

Peirce, J., Gray, J.R., Simpson, S., MacAskill, M., Höchenberger, R., Sogo, H., Kastman, E., Lindeløv, J.K., 2019. PsychoPy2: Experiments in behavior made easy. Behavior Research Methods 51, 195–203. https://doi.org/10.3758/s13428-018-01193-y

Pernet, C.R., Appelhoff, S., Gorgolewski, K.J., Flandin, G., Phillips, C., Delorme, A., Oostenveld, R., 2019. EEG-BIDS, an extension to the brain imaging data structure for electroencephalography. Scientific Data 6, 103. https://doi.org/10.1038/s41597-019-0104-8

Pion-Tonachini, L., Kreutz-Delgado, K., Makeig, S., 2019. ICLabel: An automated electroencephalographic independent component classifier, dataset, and website. NeuroImage 198, 181–197. https://doi.org/10.1016/j.neuroimage.2019.05.026

Powell, L.J., Kosakowski, H.L., Saxe, R., 2018. Social Origins of Cortical Face Areas. Trends in cognitive sciences 22, 752–763. https://doi.org/10.1016/J.TICS.2018.06.009

Ritchie, J.B., Zeman, A.A., Bosmans, J., Sun, S., Verhaegen, K., Op de Beeck, H.P., 2021. Untangling the Animacy Organization of Occipitotemporal Cortex. Journal of Neuroscience 41, 7103–7119. https://doi.org/10.1523/JNEUROSCI.2628-20.2021

Seijdel, N., Loke, J., van de Klundert, R., van der Meer, M., Quispel, E., van Gaal, S., de Haan, E.H.F., Scholte, H.S., 2021. On the Necessity of Recurrent Processing during Object Recognition: It Depends on the Need for Scene Segmentation. Journal of Neuroscience 41, 6281–6289. https://doi.org/10.1523/JNEUROSCI.2851-20.2021

Sliwa, J., Freiwald, W.A., 2017. A dedicated network for social interaction processing in the primate brain. Science 356, 745–749. https://doi.org/10.1126/science.aam6383

Smith, S.M., Nichols, T.E., 2009. Threshold-free cluster enhancement: addressing problems of smoothing, threshold dependence and localisation in cluster inference. NeuroImage 44, 83–98. https://doi.org/10.1016/J.NEUROIMAGE.2008.03.061

Tang, H., Schrimpf, M., Lotter, W., Moerman, C., Paredes, A., Caro, J.O., Hardesty, W., Cox, D., Kreiman, G., 2018. Recurrent computations for visual pattern completion. Proceedings of the National Academy of Sciences of the United States of America 115, 8835–8840. https://doi.org/10.1073/PNAS.1719397115

Wardle, S.G., Taubert, J., Teichmann, L., Baker, C.I., 2020. Rapid and dynamic processing of face pareidolia in the human brain. Nat Commun 11, 4518. https://doi.org/10.1038/s41467-020-18325-8

Yargholi, E., Op de Beeck, H.P., 2022. Category trumps shape as an organizational principle of object space in the human occipitotemporal cortex. bioRxiv 2022.10.19.512675. https://doi.org/10.1101/2022.10.19.512675

Yilmaz, M., Meister, M., 2013. Rapid innate defensive responses of mice to looming visual stimuli. Current Biology 23, 2011–2015. https://doi.org/10.1016/J.CUB.2013.08.015

